# Compartmentalized Cell Envelope Biosynthesis in *Mycobacterium tuberculosis*

**DOI:** 10.1101/2022.01.07.475471

**Authors:** Julia Puffal, Ian L. Sparks, James R. Brenner, Xuni Li, John D. Leszyk, Jennifer M. Hayashi, Scott A. Shaffer, Yasu S. Morita

## Abstract

The intracellular membrane domain (IMD) is a metabolically active and laterally discrete membrane domain initially discovered in *Mycobacterium smegmatis*. The IMD correlates both temporally and spatially with the polar cell envelope elongation in *M. smegmatis*. Whether or not a similar membrane domain exists in pathogenic species remains unknown. Here we show that the IMD is a conserved membrane structure found in *Mycobacterium tuberculosis*. We used two independent approaches, density gradient fractionation of membrane domains and visualization of IMD-associated proteins through fluorescence microscopy, to determine the characteristics of the plasma membrane compartmentalization in *M. tuberculosis*. Proteomic analysis revealed that the IMD is enriched in metabolic enzymes that are involved in the synthesis of conserved cell envelope components such as peptidoglycan, arabinogalactan, and phosphatidylinositol mannosides. Using a fluorescent protein fusion of IMD-associated proteins, we demonstrated that this domain is concentrated in the polar region of the rod-shaped cells, where active cell envelope biosynthesis is taking place. Proteomic analysis further revealed the enrichment of enzymes involved in synthesis of phthiocerol dimycocerosates and phenolic glycolipids in the IMD. We validated the IMD association of two enzymes, α1,3-fucosyltransferase and fucosyl 4- *O*-methyltransferase, which are involved in the final maturation steps of phenolic glycolipid biosynthesis. Taken together, these data indicate that functional compartmentalization of membrane is an evolutionarily conserved feature found in both *M. tuberculosis* and *M. smegmatis*, and *M. tuberculosis* utilizes this membrane location for the synthesis of its surface-exposed lipid virulence factors.

**IMPORTANCE:** *M. tuberculosis* remains an important public health threat, with more than one million deaths every year. The pathogen’s ability to survive in the human host for decades highlights the importance of understanding how this bacterium regulates and coordinates its metabolism, cell envelope elongation, and growth. The IMD is a membrane structure that associates with the subpolar growth zone of actively growing mycobacteria, but its existence is only known in a non-pathogenic model, *M. smegmatis*. Here, we demonstrated the presence of the IMD in *M. tuberculosis*, making the IMD an evolutionarily conserved plasma membrane compartment in mycobacteria. Furthermore, our study revealed that the IMD is the factory for synthesizing phenolic glycolipids, virulence factors produced by slow-growing pathogenic species.

## INTRODUCTION

Mycobacteria grow by depositing *de novo* synthesized cell envelope materials near the pole rather than the main sidewall of the rod-shaped cell (1–5). The cell envelope of mycobacteria is complex, composed of plasma membrane, peptidoglycan, arabinogalactan, and mycomembrane (6–9), making the spatially restricted cell envelope elongation a unique challenge for these bacteria. Precise coordination of the synthesis of cell envelope components are likely crucial, as highlighted by numerous proteins localizing to specific subcellular regions in mycobacteria (10). Among them, DivIVA, a filamentous coiled-coil protein, is associated with the pole of the cell and its physical interaction with early cell elongation machinery is implicated in the determination of the site of cell envelope elongation (11, 12). The recruitment of other enzymes involved in cell envelope biosynthesis defines a subpolar growth zone where new envelope components are made and incorporated into the growing pole (5, 11, 13). Polar enrichment is observed not only for proteins but also for membrane lipids. Plasma membrane heterogeneity was first reported using fluorescent lipid probes (14), and a more recent work suggested an enrichment of cardiolipin to the pole of growing mycobacterial cells (15).

Previously, we reported the presence of a growth-pole associated membrane domain, the intracellular membrane domain (IMD), in the nonpathogenic model organism *Mycobacterium smegmatis* (16–18). This domain is separated from the conventional plasma membrane upon density gradient fractionation of mycobacterial cell lysate. Plasma membrane is copurified with the cell wall, apparently due to physical connections between the two (termed PM-CW). The IMD, in contrast, is purified as ∼50 nm membrane vesicles without the association of cell wall components. Among the proteins associated with the IMD, we find essential enzymes involved in lipid biosynthetic reactions. For instance, phosphatidylinositol mannosides (PIMs) are a major component of mycobacterial plasma membrane, and the initial phase of its biosynthesis is enriched in the IMD (16, 17). The IMD is not only present for cell envelope biosynthesis but also for other membrane-associated metabolisms. Menaquinone is a major respiratory chain electron carrier in mycobacteria, and the final maturation steps of its biosynthesis take place in the IMD (19). These observations indicate the diversity of crucial processes required for polar cell envelope growth and central metabolism that are associated with this membrane domain.

In *Mycobacterium tuberculosis*, we know comparatively less about subcellular localizations of membrane proteins. DivIVA is associated to the poles in *M. tuberculosis* (11) and CwsA, which plays a role in cell wall synthesis and interacts with DivIVA, also shows polar and mid-cell localization (20). On the other hand, FtsZ and its interacting partner SepF localize to future division sites (21, 22). Furthermore, the sensor kinase MtrB localizes to septa in *M. tuberculosis* as well (23). These past studies show conserved features of pole- and septum-associated membrane proteins. However, other types of membrane compartmentalization have not been reported in pathogenic mycobacteria. In this study, we investigated the presence of the IMD in *M. tuberculosis*. Using subcellular fractionation, we first examined the IMD proteome in comparison to the PM-CW proteome. We then visualized the localization of the IMD in live growing *M. tuberculosis*. Our results demonstrate that the IMD is a membrane domain conserved in mycobacteria.

## RESULTS

### Density gradient fractionation of the *M*. *tuberculosis* cell lysate

We prepared a lysate of exponentially growing *M*. *tuberculosis* cells and fractionated by density gradient sedimentation (**Fig. 1A**). As previously seen in *M*. *smegmatis*, the majority of proteins were enriched in the top two fractions. In the density gradient of *M. smegmatis* lysate, cytoplasmic proteins are retained in the top fractions and do not sediment significantly. To confirm that the *M. tuberculosis* proteins in the top fractions correspond to cytoplasmic proteins, we performed immunoblotting against the cytoplasmic enzyme Ino1, an inositol-3-phosphate synthetase (24), and detected this protein predominantly in fractions 1 - 2 (**Fig. 1B**). Next, we determined the localization of the mannosyltransferase MptC, a polytopic membrane protein found in the PM-CW in *M. smegmatis*, by immunoblotting. MptC was enriched in the denser region of the gradient, spanning from fractions 8 - 11 (**Fig. 1B**). These fractions correspond to the density of 1.127-1.154 g/ml, which is comparable to 1.131-1.159 g/ml for *M. smegmatis* PM-CW (19), suggesting that MptC is a PM-CW protein in *M. tuberculosis* as well.

**Figure 1.**
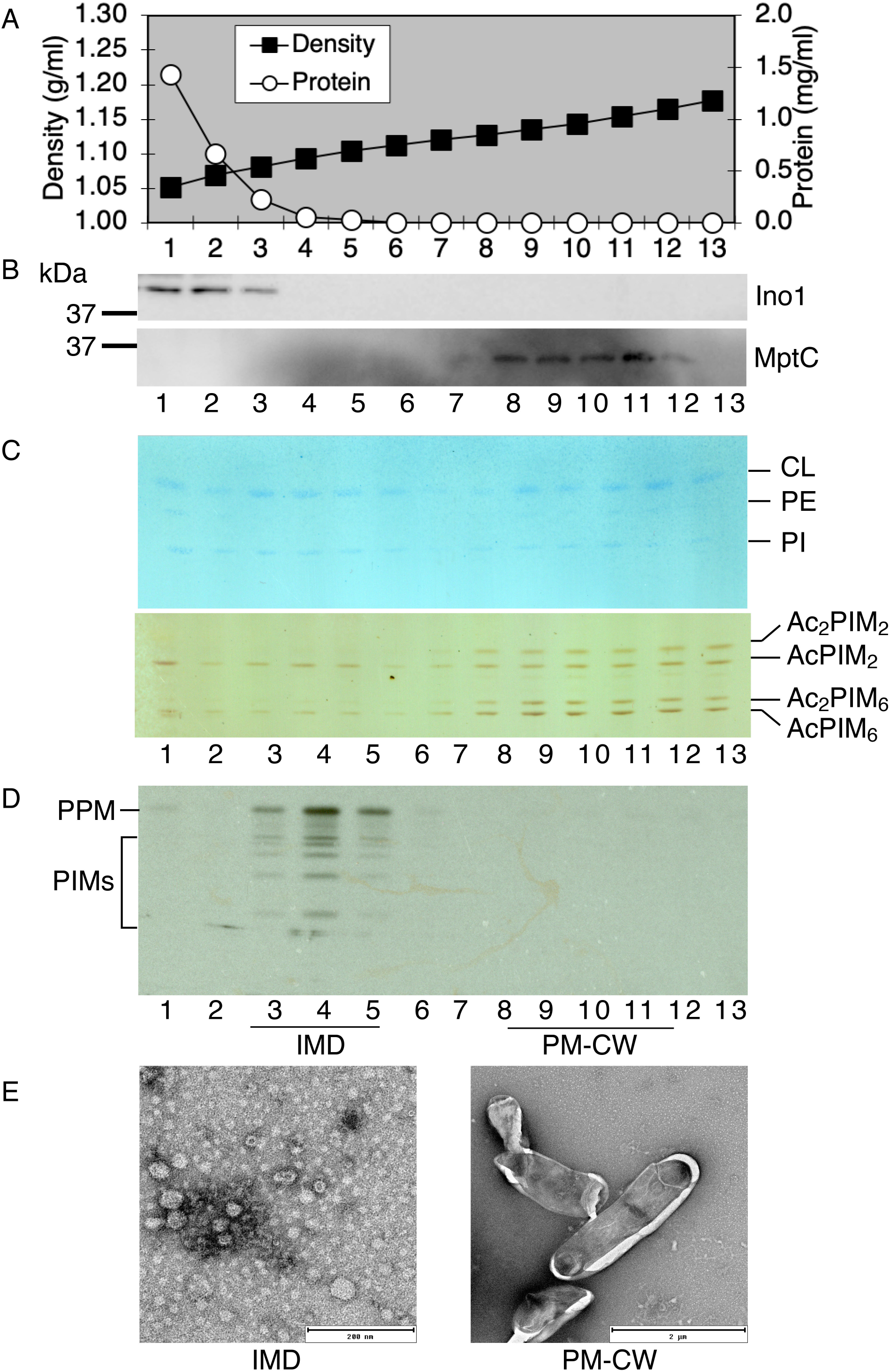
Membrane compartmentalization in *M. tuberculosis* mc^2^6230. **A**. Sucrose density gradient fractionation of attenuated *M. tuberculosis* strain. The sucrose density and protein concentration of the fractions are shown. **B**. Western blot detection of cytoplasmic marker Ino1 (40 kDa) and PM-CW marker MptC (47 kDa). **C**. Distribution of membrane phospholipids across the gradient fractions. Phospholipids were stained by molybdenum blue staining. PIMs were detected by orcinol staining. **D.** Cell-free PPM biosynthesis detection by GDP-[^3^H]-Mannose radiolabeling. **F.** Negative staining EM of the IMD and PM-CW fractions.

To examine if the IMD exists in *M. tuberculosis* in addition to the PM-CW fraction, we examined the compositional profiles of plasma membrane phospholipids. Lipids were extracted from each fraction and visualized by chemical staining. As shown in **Fig. 1C**, membrane phospholipids cardiolipin (CL), phosphatidylethanolamine (PE), phosphatidylinositol (PI) and PIMs were detected in the PM-CW fractions as well as in another density region spanning from fractions 3 to 6. These data suggest that there is an additional membrane fraction separate from the PM-CW. One known reaction that takes place in *M. smegmatis* IMD is the biosynthesis of polyprenol-phosphate-mannose (PPM). We have previously shown that the PPM biosynthetic activity is enriched in the IMD and the enzyme Ppm1 was found exclusively in the IMD in *M*. *smegmatis* (16, 17). To determine the localization of this biosynthetic reaction in *M*. *tuberculosis*, we performed a cell-free radiolabeling assay using GDP-[^3^H]mannose as a mannose donor. As shown previously (25–27), *M. smegmatis* produces heptapentenyl and decapentenyl PPM species (**Fig. S1**). In contrast, *M. tuberculosis* does not produce heptapentenyl polyprenols, and decapentenyl PPM is the major product (27–29). Although the enzyme activity is not as robust as that of *M. smegmatis* enzyme, we detected the synthesis of the expected decapentenyl PPM from *M. tuberculosis* cell lysate (**Fig. S1**), and was enriched in fractions 3 - 5 of a sucrose density gradient (**Fig. 1D**), suggesting the subcellular localization of PPM synthetase in these membrane fractions.

To compare the morphology of the membrane present in fractions 3 - 5 with that of the PM-CW, we precipitated these membrane fractions by differential centrifugation and visualized by negative staining electron microscopy (EM) (**Fig. 1E**). As in *M*. *smegmatis* (17), we observed small vesicle-like structures in fractions 3 - 5, while the PM-CW fraction revealed larger fragments of cells. Together, these data support that fractions 3 - 5 correspond to the IMD in *M*. *tuberculosis* and its overall characteristics are similar to those of the *M. smegmatis* IMD.

### Identification of IMD-associated proteins in *M. tuberculosis*

To uncover the IMD proteome, we purified the membranes in both IMD and PM-CW fractions by differential centrifugation and analyzed the trypsin digests of total proteins by liquid chromatography-mass spectrometry (LC-MS) in biological triplicates. The comparative proteomes of the two membranes were distinct with a total of 64 or 298 proteins enriched more than 2-fold (p < 0.05) in the PM-CW or IMD, respectively (**Fig. 2A**, **Dataset S1**). PM-CW was associated with enzymes with activities that are related to the typical functions of plasma membrane (**Fig. 2B**). For example, we identified Esx5 type VII secretion system components EccB5-E5 (Rv1782, Rv1783, Rv1795), trehalose monomycolate transporter MmpL3 (Rv0206c), a subunit of the phosphate specific transport system, PstS1 (Rv0934), and NADH-quinone oxidoreductase subunits NuoF and NuoG (Rv3150, Rv3151). The IMD, on the other hand, was more restricted to enzymes involved in lipid metabolism (**Fig 2B**). For instance, we found enzymes involved in the synthesis of polyprenol-linked galactan precursor for arabinogalactan synthesis enriched in the IMD: the GlcNAc-diphospho-decaprenol L-rhamnosyltransferase WbbL1 (Rv3265c), the Rha-GlcNAc-diphospho-decaprenol β1,4/1,5-galactofuranosyltransferase GlfT1 (Rv3782) and the Gal*f*-Gal*f*-Rha-GlcNAc-diphospho- decaprenol β1,5/1,6-galactofuranosyltransferase GlfT2 (Rv3808c). The PPM synthase Ppm1 (Rv2051c) and acyl-CoA dehydrogenases such as FadE24 (Rv3139), FadE23 (Rv3140), and FadE10 (Rv0873) were also found in the IMD. From the total of 298 proteins identified in the IMD through the comparative proteomics, 218 had *M*. *smegmatis* homologues. Of these, 101 have also been identified previously as enriched in the IMD in a comparative proteomics of *M*. *smegmatis* IMD and PM-CW (**Dataset S1**) (16). These data are consistent with the idea that the *M. tuberculosis* IMD is a metabolically active membrane domain with multiple functions shared with that of *M. smegmatis* (see below for a more refined IMD proteome).

**Figure 2.**
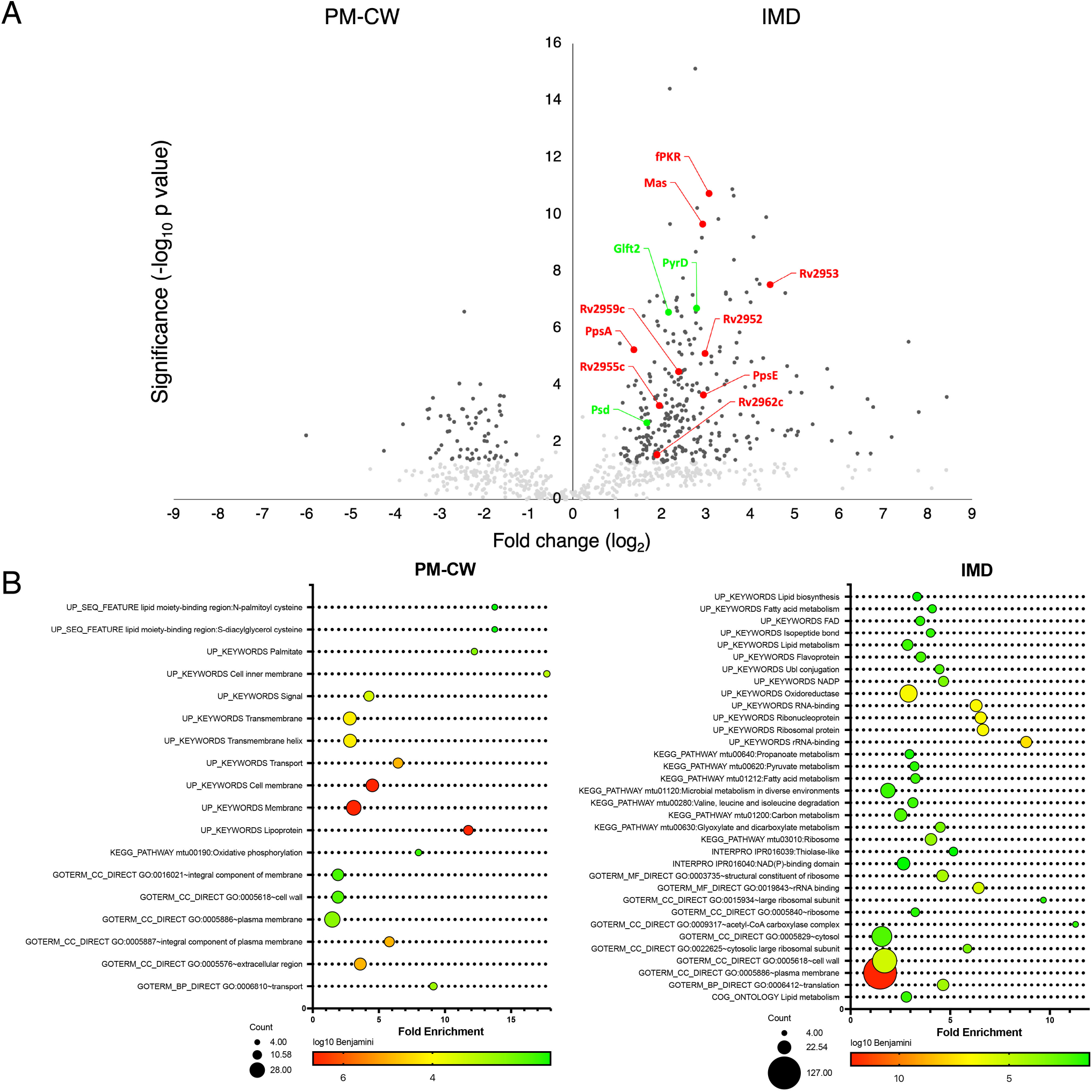
Comparative proteome of the conventional plasma membrane (PM-CW) and the IMD. Cell growth, lysate preparation, membrane purification and proteomic analysis were repeated three times. **A.** The volcano plot of proteins enriched in either the PM-CW or the IMD. Proteins that were enriched more than two-fold with p < 0.05 were considered significant (black dots). Other identified proteins are shown in gray dots. Red dots correspond to significantly enriched proteins involved in PGL/PDIM biosynthesis. Green dots correspond to other proteins analyzed biochemically and/or by fluorescent microscopy in this study. **B.** Functional annotation of proteins enriched in the membrane fractions using UniProt keywords, showing that the IMD is enriched in proteins related to membrane and lipid metabolism. The enrichment represents the proportion of the term in the IMD proteome compared to its expected proportion in the genome. The area of the circles are proportional to the number of proteins in each category.

### Validation of IMD-associated proteins

Among the IMD-associated proteins was the dihydroorotate dehydrogenase PyrD, which is involved in pyrimidine biosynthesis and is dependent on a lipidic electron carrier, menaquinone, for the redox reaction (30, 31). Because it is an IMD-associated protein in *M. smegmatis* (16), we chose this protein for the validation of the *M. tuberculosis* IMD proteome. We cloned the gene with a hemagglutinin (HA) epitope tag into an expression vector, integrated into the bacteriophage L5 integration site of *M. tuberculosis* genome, and confirmed its expected molecular weight of 42.1 kDa (**Fig. 3A**). We then fractionated the cell lysate by sucrose density gradient (**Fig. 3B**), and confirmed the IMD localization of PyrD-HA in contrast to the PM-CW localization of MptC (**Fig. 3C**). These data indicate that PyrD associates with the IMD in *M. tuberculosis* validating the comparative proteomics analysis.

**Figure 3.**
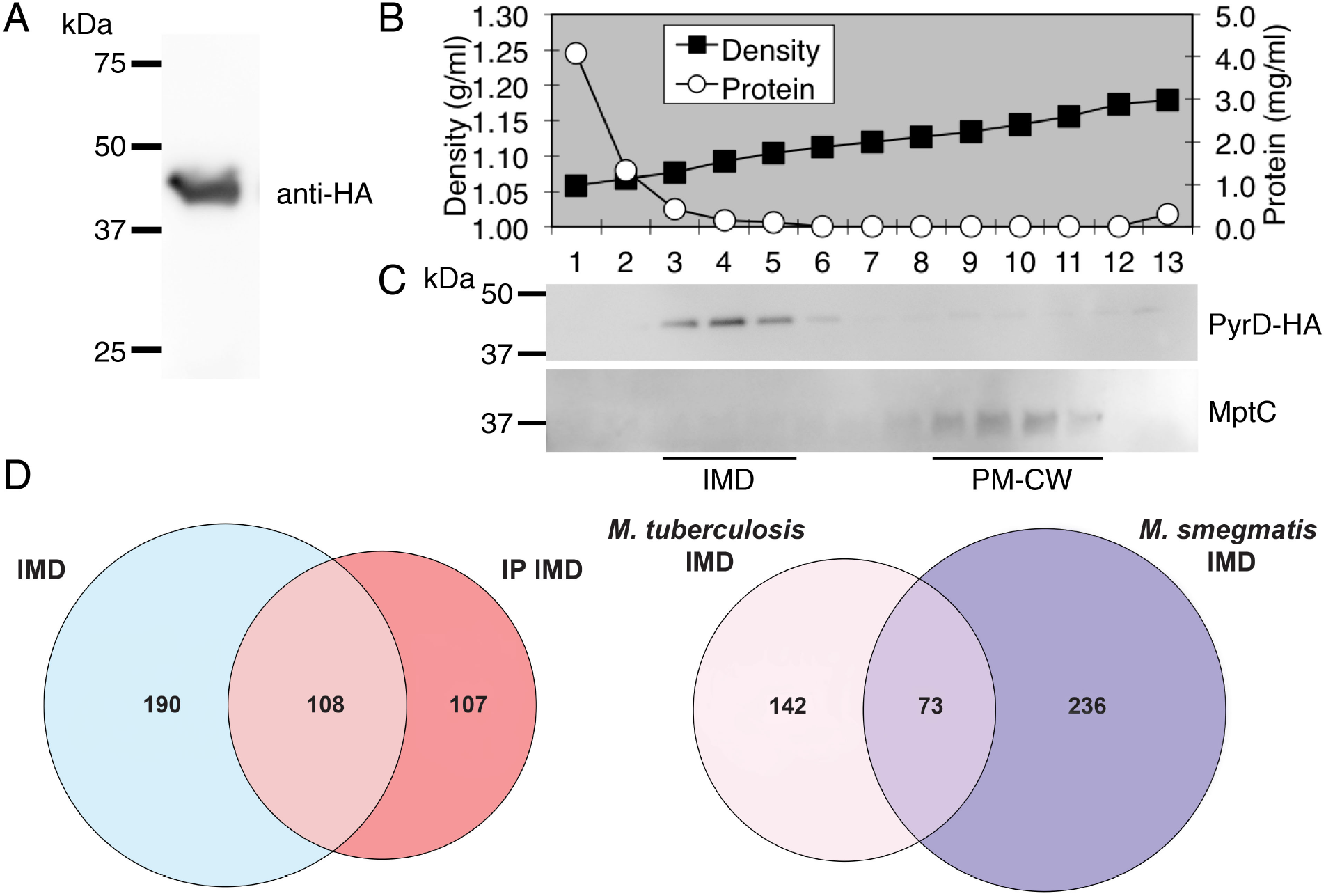
PyrD is an IMD-associated protein. **A**. Western blotting of a crude lysate prepared from a merodiploid *M. tuberculosis* strain producing PyrD-HA. In this strain, the expression vector was integrated at L5 *attB* site. Expected molecular weight of PyrD-HA is 42.1 kDa. **B.** Sucrose density gradient fractionation. The density and protein concentration for each fraction are shown. **C**. Western blot detection of PyrD-HA in the IMD fractions and the PM-CW marker MptC (Rv2181, 47 kDa). **D.** Proteome of an immunoprecipitated IMD. IMD was purified from the PyrD-HA-expressing strain using anti-HA antibody. Left, Venn diagram showing the overlap of proteins found in the comparative IMD proteome (Fig. 2) and immunoprecipitated IMD proteome. Right, Venn diagram showing the orthologs found commonly in the immunoprecipitated IMD proteomes of *M. tuberculosis* and *M. smegmatis*.

Next, we took advantage of the PyrD-HA-expressing strain and purified the IMD from the gradient fractions by vesicle immunoprecipitation using anti-HA agarose beads as performed previously in *M. smegmatis* (16). We used the IMD fraction from the parental strain as a negative control for immunoprecipitation. With a cutoff of 5-fold enrichment between the experimental and negative control samples, 215 proteins were identified in the IMD of *M*. *tuberculosis* (**Dataset S1**). Among them, 108 proteins were found in the IMD proteome of the initial comparative proteomics (**Fig. 3D**). Out of these 215 immunoprecipitated proteins, 148 had *M*. *smegmatis* homologues, with 73 localizing in the IMD of both species (**Fig. 3D**). Sixty seven IMD-associated proteins were unique to *M*. *tuberculosis*, and notable proteins among them were enzymes involved in the biosynthesis of phthiocerol dimycocerosate (PDIM) and phenolic glycolipids (PGL; see below for further analysis). These data indicate that the IMD is an evolutionarily conserved domain in mycobacteria with species-specific features.

### Visualization of IMD proteins in *M. tuberculosis*

To visualize the IMD in *M. tuberculosis*, we constructed a mutant, in which a gene encoding 2xHA-tagRFP tag was knocked into the 3’ end of the endogenous *pyrD* gene locus. The in-frame insertion was designed to produce a PyrD-2xHA-tagRFP fusion protein from the endogenous locus using the native promoter. This mutant was grown to log phase, and cell lysate was fractionated by sucrose density gradient. We detected the localization of this fusion protein to the IMD by western blotting using anti-HA antibody (**Fig. 4**). When we examined this strain by live fluorescence microscopy, we could not detect the fluorescence (not shown), possibly due to low abundance of the protein expressed from the endogenous promoter.

**Figure 4.**
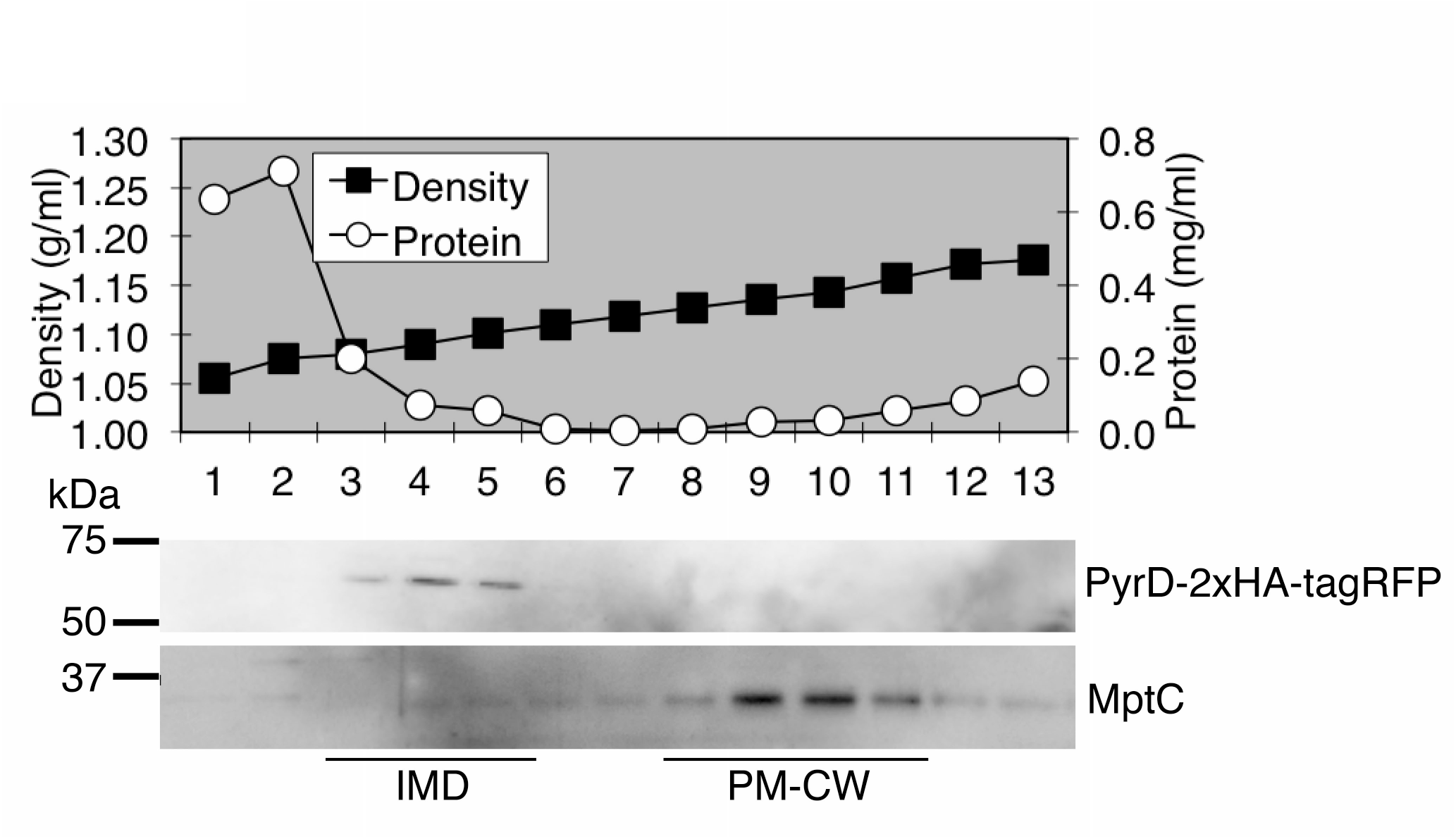
PyrD-2xHA-tagRFP, expressed from the endogenous locus, showing IMD association. **A**. Sucrose gradient density fractionation of *M. tuberculosis* strain where the endogenous *pyrD* gene was replaced with *pyrD-2xHA-tagRFP*. Sucrose density and protein concentration for each fraction are shown. **B**. Western blot showing the IMD localization of PyrD-2xHA-tagRFP (68.2 kDa). MptC, PM-CW marker (47 kDa). Experiments are repeated twice and representative results are shown.

As an alternative approach, we next used an expression vector that integrates at the mycobacteriophage L5 *attB* site and expressed an IMD-associated protein as fluorescent protein fusions using a strong promoter. We chose two proteins: GlfT2, a galactosyltransferase involved in galactan biosynthesis, and Psd, a phosphatidylserine decarboxylase involved in PE biosynthesis (see **Fig. 2A**). These enzymes are also enriched in the IMD of *M*. *smegmatis* (16, 17). We fractionated the lysates of cells expressing either GlfT2-mNeonGreen-HA or Psd- mNeonGreen-HA. Both GlfT2-mNeonGreen-HA and Psd-mNeonGreen-HA showed expected molecular weights (100.9 and 53.5 kDa, respectively), and were enriched in the IMD fractions (**Fig. 5A-B and S2A-B**) although we noted that there was a weak signal of Psd-mNeonGreen-HA in the PM-CW.

**Figure 5.**
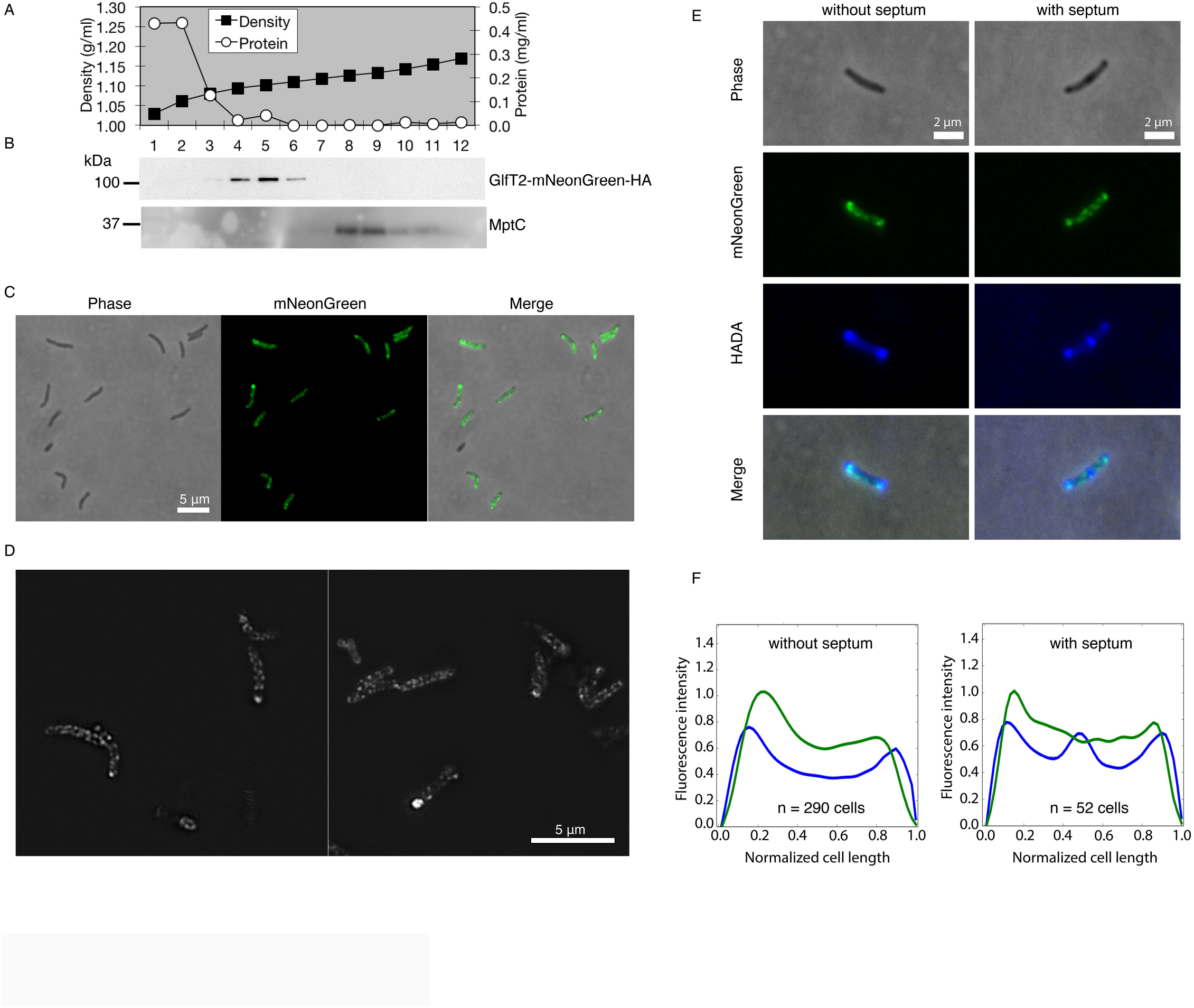
Polar enrichment and sidewall patches of GlfT2-mNeonGreen-HA, an IMD-associated protein. **A.** Density gradient and protein concentration profile. **B.** Western blotting of GlfT2- mNeonGreen-HA (100.9 kDa). MptC, PM-CW marker. **C.** Conventional fluorescence microscopy images of log phase cells. **D.** SIM images of log phase cells. **E.** Cells expressing GlfT2-mNeonGreen-HA, labeled with HADA for 2 hours and visualized by conventional fluorescence microscopy. **F.** Fluorescence intensity profile of GlfT2-mNeonGreen-HA from cells labeled with HADA, for which representative images are shown in the panel E. Cells were categorized into two populations based on the presence or absence of a septum, as determined by HADA labeling. Intensity profiles are aligned with the brightest pole on the left and are shown in arbitrary units. Green and blue fluorescence intensity profiles correspond to GlfT2- mNeonGreen-HA and HADA respectively. Experiments are repeated at least twice and representative results are shown.

We then analyzed the localization of these proteins in live cells by fluorescence microscopy. In a subset of cells, GlfT2-mNeonGreen-HA showed intense fluorescence near the poles of the cells with less intense patches of fluorescence along the sidewall membrane (**Fig. 5C**). This fluorescence pattern is consistent with the IMD fluorescence patterns observed in *M*. *smegmatis*. We used structured illumination microscopy (SIM) to acquire higher-resolution images. As shown in **Fig. 5D**, intense foci near the poles of the cells were evident and the sidewall patches appeared consistent with the idea that the protein is a membrane-associated protein. Even though the polar enrichment of GlfT2-mNeonGreen-HA was evident in some cells, we often found many cells where the intensities of polar and sidewall patches were similar, making the polar enrichment not as obvious as in the case of *M. smegmatis*. We considered the possibility that the polar IMD enrichment may correlate with a specific cell cycle stage. To visualize cells with or without an active septum synthesis, we labeled the cells with 3-[[(7-hydroxy-2-oxo-2H-1-benzopyran-3-yl)carbonyl]amino]-D-alanine (HADA), a fluorescent D-amino acid analog (**Fig. 5E**), and separated the cell population based on the presence of a septum.

We then determined the average fluorescence intensity profiles of GlfT2-mNeonGreen-HA along the cell length as we have previously done in *M. smegmatis* (32). We observed that the enrichment of GlfT2-mNeonGreen-HA was slightly proximal to the polar HADA labeling, and was evident regardless of whether there is an active septum synthesis or not (**Fig. 5F**).

Comparable fluorescence patterns were also observed for Psd-mNeonGreen-HA (**Fig. S2C**). Together, these data show that the IMD is a membrane domain that is frequently enriched in the subpolar region of the cell.

### Maturation of PGL takes place in the IMD

The biosynthesis pathways of PDIM and PGL are partially overlapping, utilizing the same enzymes for several steps (**Fig. 6A**). The pathway can be divided into synthesis, assembly, and maturation stages. In the synthesis stage, mycocerosates and (phenol)phthiocerols are synthesized by Mas and PpsA-E, respectively. In the assembly stage, PapA5 esterifies two molecules of mycocerosates to the hydroxyl groups of phthiocerol or phenolphthiocerol, producing PDIM or a PGL precursor, phenolphthiocerol dimycocerosate (PPDIM), respectively. In the maturation stage, PPDIM is further decorated at the hydroxyl group of the phenol residue by several carbohydrate residues and these carbohydrates are modified by methylation to become a mature PGL.

**Figure 6.**
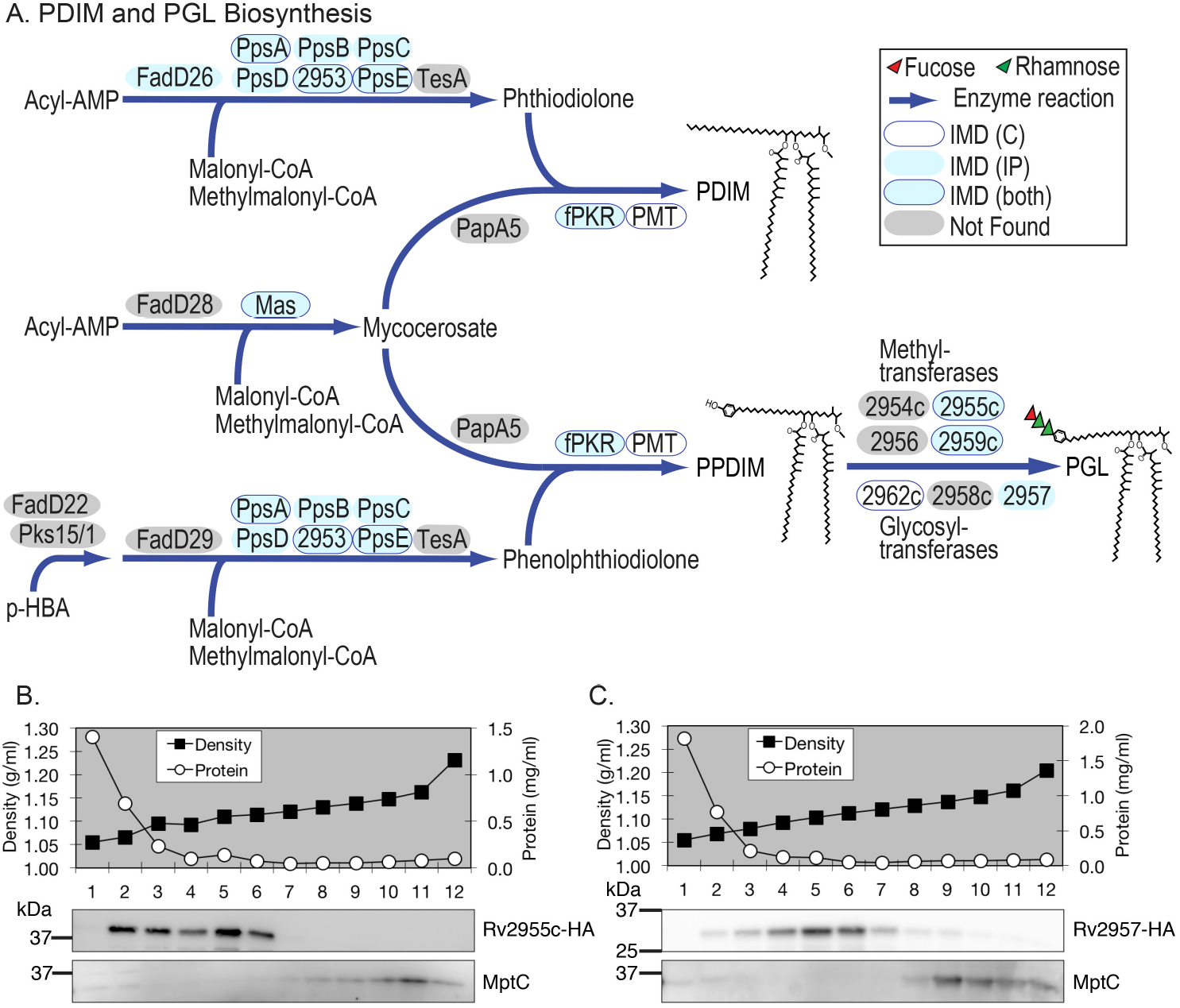
PDIM and PGL biosyntheses take place in the IMD. **A.** Biosynthetic pathways of PDIM and PGL. Enzymes involved in the biosyntheses are shown in ovals, which are color-coded to indicate the subcellular localization based on proteomic analyses. **B.** Western blotting of Rv2955c-HA (37.8 kDa). **C.** Western blotting of Rv2957-HA (32.9 kDa). MptC, PM-CW marker.

We found several enzymes in these pathways enriched in the IMD (**Fig. 6A**). Among them, we generated merodiploid strains producing HA epitope-tagged PpsB, PpsE, and fPKR (Rv2951c). PpsB and PpsE are polyketide synthases involved in the synthesis of (phenol)phthiocerols while fPKR is a F_420_H_2_-dependent phthiodiolone ketoreductase (33–35). PpsE and fPKR were enriched in the IMD based on the comparative analysis of IMD / PM-CW proteomes as well as the proteome of immunoprecipitated IMD. PpsB was not found in the comparative proteome, but identified in the immunoprecipitated IMD. We analyzed lysates of the merodiploid strains by SDS-PAGE and anti-HA immunoblot, but were unable to detect any of these epitope-tagged proteins (not shown). Next, we generated merodiploid strains producing mNeonGreen-HA-tagged enzymes involved in the final maturation steps of PGL synthesis, namely a rhamnosyl *O*- methyltransferase (Rv2959c), a fucosyltransferase (Rv2957), and a fucosyl *O*-methyltransferase (Rv2955c). All three enzymes were enriched in the immunoprecipitated IMD. Rv2955c and Rv2959c were also enriched in the IMD based on the IMD / PM-CW comparison. When mNeonGreen-HA-tagged proteins were analyzed by immunoblot, we found significant amounts of protein bands that were smaller than the expected molecular weight (not shown). We suspect that those fluorescent protein-tagged fusions may be prone to protein degradation. To circumvent the problem, we generated merodiploid strains producing fusion proteins tagged with a short HA epitope alone. Among them, we detected a protein band that migrated near the expected molecular weight for Rv2955c and Rv2957 whereas we could not detect Rv2959c at a significant level. We fractionated a cell lysate from the strain producing either Rv2955c-HA or Rv2957-HA, and determined the subcellular localization. Both proteins were localized to the IMD although Rv2955c was distributing more broadly to a lighter density region as well (**Fig. 6B and C**). Thus, our biochemical analysis validates that the final maturation steps of PGL synthesis takes place in the IMD. Taken together with proteomic identifications, out data support the idea that the majority of enzymatic reactions to synthesize PDIM/PGL take place in the IMD.

## DISCUSSION

The IMD is proposed to provide a membrane surface for the cell envelope biosynthesis and growth of *M. smegmatis*, a non-pathogenic model *Mycobacterium* (16, 17). Here, we demonstrated that a similar membrane organization exists in *M*. *tuberculosis*. Furthermore, our data demonstrate that the biosynthesis of PGL, a virulence factor produced by a subset of pathogenic mycobacteria (36–40), takes place in the IMD. We suggest that the IMD is an evolutionary conserved subcellular organization in mycobacteria.

A number of metabolic reactions in the IMD are conserved between the two species. First, GlfT2, a galactofuranosyltransferase involved in arabinogalactan synthesis, was found in the IMD of both species. Two reactions preceding the GlfT2-driven galactose chain elongation are mediated by the rhamnosyltransferase WbbL1 (Rv3265c) and the galactofuranosyltransferase GlfT1 (Rv3782). Both WbbL1 and GlfT1 are enriched in the IMD proteome of *M. smegmatis* as well as *M. tuberculosis*, indicating that galactan precursor biosynthesis is a conserved metabolic function of the IMD. Second, polyprenol-phosphate-mannose biosynthesis is IMD-associated in *M. smegmatis* and the enzyme Ppm1 is enriched in this membrane domain (16, 17). In *M. tuberculosis*, we used radiolabeled GDP-mannose to show the enzyme activity of Ppm1 in the IMD. Furthermore, we detected Ppm1 (Rv2051c) in the *M. tuberculosis* comparative IMD proteome although we could not detect this protein from the immunoprecipitated IMD. In *M. smegmatis*, the PPM synthase Ppm1 and the integral membrane protein Ppm2 are produced separately, while they are produced as a single fused polypeptide in *M. tuberculosis* (41). The larger and more hydrophobic nature of *M. tuberculosis* Ppm1 may have contributed to the inefficient detection of this protein by proteomics. Third, PyrD, a menaquinone-dependent dihydroorotate dehydrogenase involved in pyrimidine biosynthesis, is an IMD-associated protein in *M. smegmatis* (16, 42), and our current study revealed it as an IMD protein in *M. tuberculosis* as well. These pathways represent some of the well-characterized evolutionarily conserved pathways enriched in the IMD. In total, our data showed that 69-73% of the IMD-associated *M. tuberculosis* proteins had *M. smegmatis* orthologs, which was comparable to the two-species overlap found in the whole genome, where 63-69% of *M. tuberculosis* protein-coding genes have *M. smegmatis* orthologs (43, 44).

It is notable that the IMD is often where final maturation steps of lipid biosynthesis take place. For example, the final step of PE biosynthesis is decarboxylation of phosphatidylserine mediated by the IMD-associated enzyme Psd (17, 42). We confirmed in this study that Psd (Rv0437c), which was expressed as a mNeonGreen-HA fusion protein, is an IMD-associated protein by both subcellular fractionation and fluorescence microscopy. Similarly, during the synthesis of an electron carrier menaquinone, demethylmenaquinone produced by the PM-CW enzyme MenA is modified by the IMD-associated demethylmenaquinone methyltransferase MenG and menaquinone reductase MenJ to become a mature form of menaquinone (19). We identified both MenG (Rv0558) and MenJ (Rv0561c) in the proteome of immunoprecipitated *M. tuberculosis* IMD, suggesting a conserved compartmentalization of these final two steps of menaquinone biosynthesis in the pathogenic species. Finally, peptidoglycan precursor lipid I matures into lipid II by a GlcNAc transfer reaction mediated by MurG, and MurG is a previously established IMD protein in *M. smegmatis* (32). In *M. tuberculosis*, MurG (Rv2153c) showed four-fold enrichment in the IMD compared to the PM-CW, although we could not detect MurG in the immunoprecipitated IMD proteome. Observations of conserved IMD-associated lipid maturation steps in multiple lipid biosynthetic pathways indicate the role of the IMD in lipid modifications.

In addition to these shared features, *M. tuberculosis* IMD revealed roles that are specific to slow-growing pathogens and not found in *M. smegmatis*. In particular, we found multiple enzymes associated with the PDIM/PGL biosynthesis (Fig. 6A). These lipids are not produced by *M. smegmatis*, and genes encoding the biosynthetic enzymes are absent in its genome. We found PpsA-E (Rv2931-2935), polyketide synthases that produce phthiodiolone and phenolphthiodiolone, enriched in the IMD proteomes. They are large proteins, ranging 159 - 231 kDa, and therefore, we were initially concerned if the IMD enrichment of PpsA and PpsE over the PM-CW might have been a consequence of these large proteins sedimenting further in a density gradient than other average-sized proteins due to their large size. However, all five enzymes were identified in the proteome of immunoprecipitated IMD but not from the mock immunoprecipitation of a wildtype cell lysate. Therefore, our data suggest that these enzymes are truly associated with the IMD. Unfortunately, our attempts to express epitope-tagged PpsB and PpsE were unsuccessful, and validation of the subcellular localization of these enzymes must await future studies. PapA5 transfers two mycocerosates to either a phthiodiolone or a phenolphthiodiolone, producing phthiodiolone dimycocerosate or phenolphthiodiolone dimycocerosate, respectively. These lipids are further modified by sequential reactions of F_420_H_2_-dependent phthiodiolone ketoreductase (fPKR, Rv2951c) and phthiotriol/phenolphthiotriol dimycocerosates methyltransferase (PMT, Rv2952) to become a mature PDIM or phenolphthiocerol dimycocerosate (PPDIM), a precursor of PGL (34, 35, 45, 46). Comparative proteomic analysis showed that both fPKR and PMT are enriched in the IMD, and fPKR was also detected in the immunoprecipitated IMD. We attempted to express fPKR with C-terminal mNeonGreen-HA or HA tag, but could not detect the protein production by immunoblotting. fPKR has a predicted molecular weight of 41.3 kDa, and has no predicted transmembrane domains. Therefore, it is likely that fPKR peripherally associates with the IMD, but further validations are needed.

Finally, we attempted to produce epitope-tagged glycosyltransferases and methyltransferases that are involved in the maturation of PPDIM to PGL. Two methyltransferases (Rv2955c and Rv2959c) and a fucosyltransferase (Rv2957) were found in the IMD proteomes. While we could not detect stable production of epitope-tagged Rv2959c, we were able to produce Rv2955c and Rv2957 with C-terminal HA tag. Both proteins were found enriched in the IMD and absent in the PM-CW, confirming the proteomic analysis. Rv2955c was more widely distributed into lighter density regions (Fractions 2-3), which is a feature observed for some *M. smegmatis* proteins as well (*e.g.* MurG, ThiD) (32, 42). The biochemically validated IMD association of two enzymes involved in the final maturation step of PGL synthesis, combined with other enzymes found in the IMD proteome, indicate that the PDIM/PGL biosynthesis takes place in the IMD.

In sum, we conclude that the IMD is a conserved membrane domain of mycobacteria enriched in lipid biosynthetic reactions.

## MATERIALS AND METHODS

### Cell cultures

*M. tuberculosis mc^2^6230 ΔRD1/panCD* strain was used throughout this study. It was grown in Middlebrook 7H9 broth or Middlebrook 7H10 agar supplemented with OADC (final concentrations: 190 µM oleic acid, 0.5% (w/v) bovine serum albumin, 11.1 mM dextrose, 13.9 mM NaCl, 0.0004% (w/v) catalase), 0.05% Tween 80, and 50 µg/ml pantothenic acid. When required, the medium was supplemented with 100 µg/ml hygromycin B (Wako), 20 µg/ml kanamycin sulfate (MP Biochemicals), 50 µg/ml streptomycin, or 5% sucrose.

### Density gradient fractionation and protein analysis

Log phase cells (OD_600_ = 0.5-1.0) were lysed and the lysate was fractionated by sucrose density gradient as described previously (16). The protein concentration for each fraction was determined by bicinchoninic acid (BCA) assay (Pierce). Sucrose density of each fraction was determined by a refractometer (ATAGO). SDS-PAGE and western blotting of the sucrose gradient fractionation were performed using equal volume of each fraction as described before (16). Anti-MptC antibody was previously raised (47) and anti-HA antibody was purchased from Sigma-Aldrich.

### Cell-free radiolabeling assay

To determine PPM biosynthesis activities, sucrose density fractions were incubated with GDP- [2-^3^H]mannose (American Radiolabeled Chemicals, 20 Ci/mmol) as described before (17), except that the time of incubation was extended to 1 hour. The lipids were extracted, purified, and resolved on high-performance thin layer chromatography (HPTLC) silica gel using chloroform/methanol/13 M ammonia/1 M ammonium acetate/water (180:140:9:9:23, v/v/v/v/v) as a solvent, and visualized by fluorography using En^3^Hance.

### Lipid analysis

Lipids were extracted from each gradient fraction as described (16) and analyzed by HPTLC as described above. Phospholipids and PIMs were detected by molybdenum blue and orcinol staining, respectively.

### Negative staining EM

Samples of pooled sucrose gradient fractions of IMD and PM-CW were pelleted at 100,000x *g* for 60 min on TLA100.2 rotor (Beckman), and washed with HES buffer (25mM HEPES-HCl, pH 7.4, 2mM EGTA, 150mM NaCl) twice. The samples were prepared for EM observation as described (16).

### Proteome preparation and analysis

Pooled IMD and PM-CW fractions were pelleted and washed, as described above, in biological triplicates. The protein samples were then separated on a 12% SDS-PAGE gel for a short distance and the gel slice containing the entire protein bands was subjected to in-gel trypsin digestion. The peptides were analyzed by nano-LC ESI MS on an Orbitrap mass spectrometer (Thermo Scientific Q Exactive) as described (16). The data were analyzed and annotated using TubercuList (Release 27, https://mycobrowser.epfl.ch/). Proteins with a minimum of two-fold enrichment (p < 0.05) in either the IMD or the PM-CW were considered to be part of the respective proteome. Functional annotation was performed using DAVID Bioinformatics Resources as described (16).

The immunoprecipitated IMD proteome was analyzed using IMD fractions from cells expressing PyrD-HA, using wild-type as a negative control. Immunoprecipitation using HA-agarose beads (Pierce), proteome analysis and functional annotation analysis were as described (16).

### Construction of plasmids

pMUM036 – To create an expression vector for PyrD (Rv2139), which is C-terminally tagged with an HA epitope, the *pyrD* gene was amplified with primers A161/A162 from *M*. *tuberculosis* genomic DNA (**Table S1**). pMUM012 (16) was digested with EcoRV and ScaI and the vector backbone was blunt-end-ligated with the PCR product.

pMUM145 – To create a knock-in strain, in which an epitope-tagged PyrD is expressed from the endogenous locus, the upstream and downstream of the *pyrD* gene were amplified using primers A536/A537 and A538/A539, respectively (**Table S1**). The PCR products were digested with Van91I. The two fragments were then ligated into Van91I-digested pCOM1 as previously described (16) to create an intermediate vector, pMUM133. Using pMV261-CO-TagRFP as a template, tagRFP was amplified and HA tag was added using primers A557/A558. The PCR product and pMUM133 were then digested with VspI and BspT1 and ligated to create a knock-in vector to insert a gene encoding 2xHA-tagRFP at the 3’ end of the *pyrD* gene.

pMUM215 – To create expression vector for C-terminally mNeonGreen-HA-tagged Psd (Rv0437c), *psd* was cloned by PCR using primers A630/A631 (**Table S1**). In a separate experiment, a gene encoding mNeonGreen was amplified from pMUM044, an unpublished plasmid which carries the same *mNeonGreen* gene as our published plasmid pMUM072 (16), using primers A483/A484. The PCR product was blunt-end-ligated with ScaI-digested pMUM098 (19) to create pMUM111. The PCR product carrying *psd* and pMUM111 were digested with NdeI and BspTI and ligated to create the final expression vector.

pMUM222 – To create an expression vector for GlfT2 (Rv3808c) with C-terminal mNeonGreen-HA epitope tag, the *glfT2* gene was amplified with primers A628/A629 (**Table S1**). The PCR product carrying *glfT2* and pMUM111 were digested with NdeI and BspTI and ligated to create the final expression vector.

pMUM257 – We first created an expression vector for Rv2955c with C-terminal mNeonGreen-HA epitope tag. The *rv2955c* gene was amplified with primers A849/850 (**Table S1**). Both the PCR product carrying *rv2955c* and pMUM235, which is a derivative of pMUM111, were digested with NdeI and BspTI and ligated to create the Rv2955c-mNeonGreen-HA expression vector (pMUM245). To remove mNeonGreen, we digested pMUM245 with PacI and AflII, blunt-ended the digested plasmid using T4 DNA polymerase, and ligated, resulting in pMUM257, the Rv2955c-HA expression vector.

pMUM258 – As in pMUM257, we first created an expression vector for Rv2957 with C-terminal mNeonGreen-HA epitope tag. The *rv2957* gene was amplified with primers A851/852 (**Table S1**). The Rv2957-HA expression vector, pMUM246, and the Rv2957-HA expression vector, pMUM258, were created following the identical procedure described for pMUM257.

Plasmid constructs were electroporated into *Mycobacterium tuberculosis mc^2^6230 ΔRD1/panCD* for integration and homologous recombination as previously described (16).

### Fluorescence microscopy

Standard fluorescence microscopy was performed as described (42). To distinguish between cells with or without septa, cells were labeled with the peptidoglycan probe HADA (Tocris Bioscience, Bristol, UK). *M. tuberculosis* cells expressing GlfT2-mNeonGreen-HA were grown to mid-log (OD_600_ = 0.5) and then labeled with 500 μM HADA for 2 hour. Labeled cells were washed twice in PBS, spotted onto a 1% agar pad slide and covered with a glass coverslip sealed to the slide with nail polish. Live cells were then imaged with a Nikon Eclipse E600 with a 100x objective lens. mNeonGreen fluorescence profiles along cell lengths were calculated with Oufti (48) and sorted based on the presence or absence of HADA-labelled septa. mNeonGreen fluorescence profiles of septated and non-septated populations were aligned by their brightest pole, normalized by cell length and then averaged using a custom Python script. For SIM, images were acquired by Nikon Eclipse Ti N-SIM E microscope equipped with a Hamamatsu Orca Flash 4.0 camera (numerical aperture,1.49) as described before (12), and reconstructed on NIS Elements imaging software.

## Acknowledgement

*M. tuberculosis mc^2^6230 ΔRD1/panCD* strain, anti-Ino1 antibody, and pMV261-CO-TagRFP plasmid were kind gifts from Drs. William Jacobs (Albert Einstein College of Medicine), Heran Darwin (New York University), and Christopher Sassetti (University of Massachusetts Medical School), respectively. Structured illumination super-resolution microscopy images were obtained at the University of Massachusetts Light Microcopy Facility (Dr. James Chambers, Director), with support from the Institute for Applied Life Sciences. This work was supported by NIH R03 AI140259 to YSM. JP was a recipient of the Science Without Borders Fellowship from CAPES-Brazil (0328-13-8).

## Supplemental Figures

**Figure S1.**
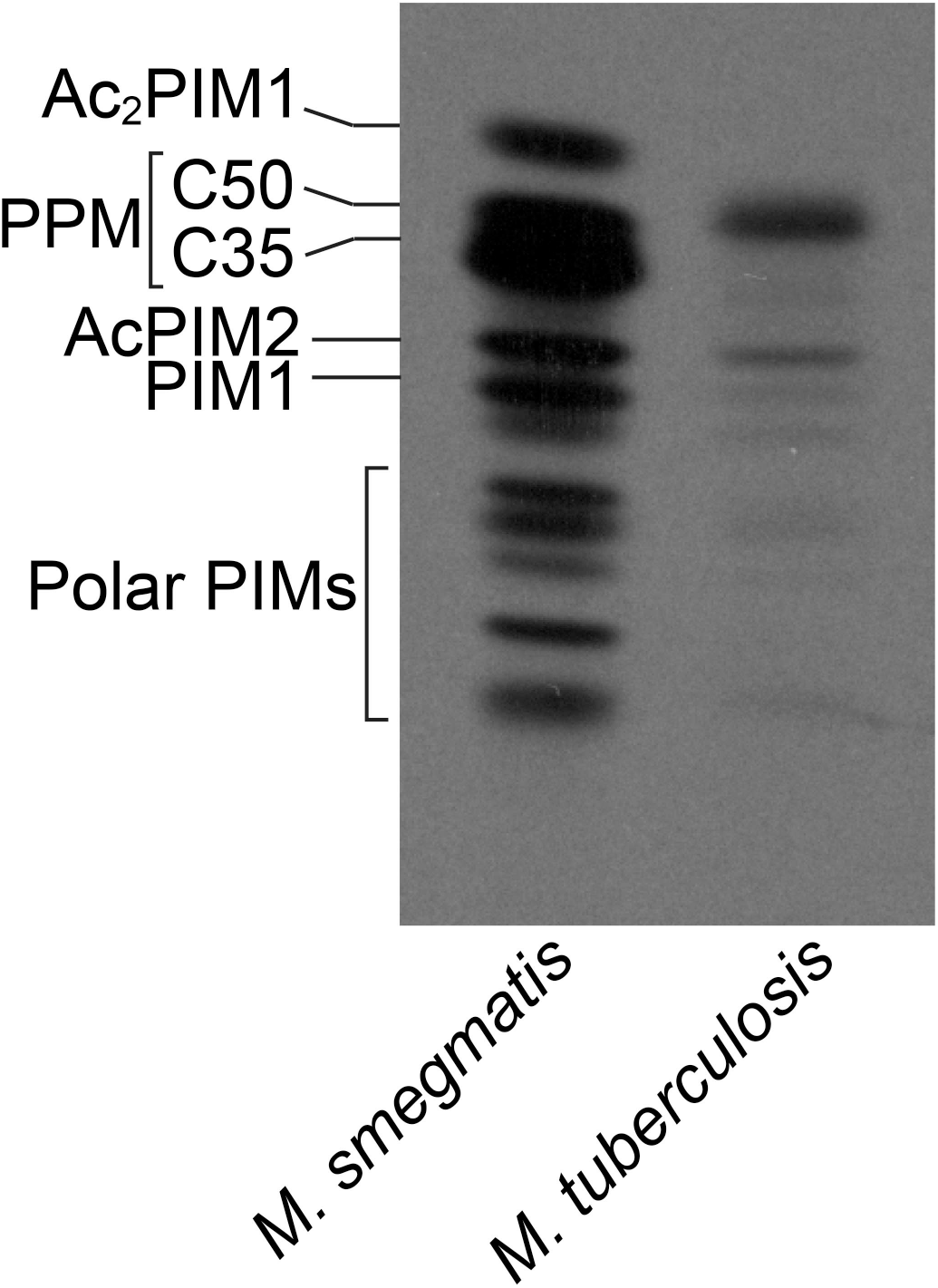
Cell-free PPM biosynthesis detection of crude lysate by GDP-[^3^H]-Mannose radiolabeling, comparing *M. smegmatis* and *M. tuberculosis*.

**Figure S2.**
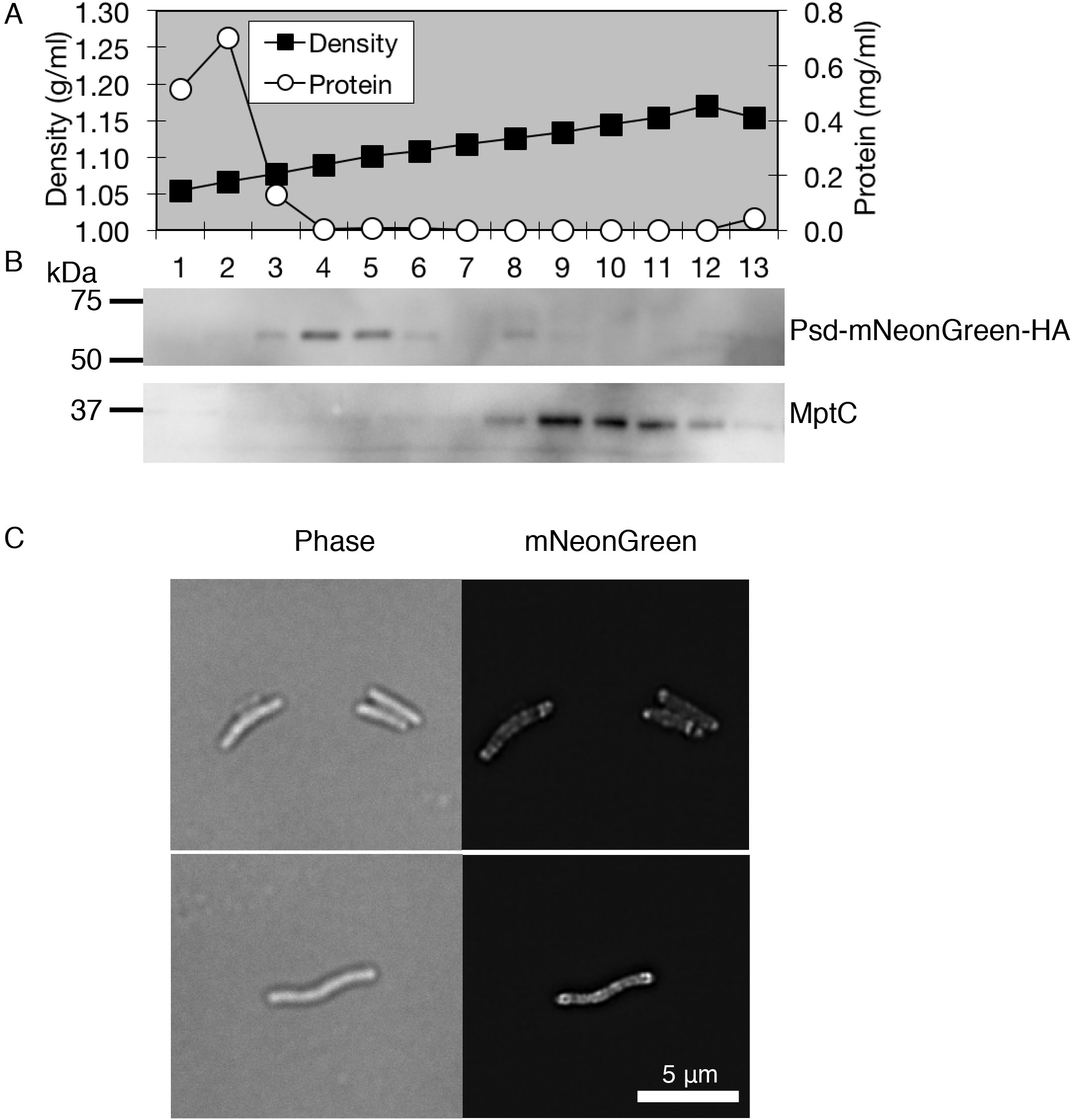
IMD association of Psd-mNeonGreen-HA. **A.** Density gradient and protein concentration profile. **B.** Western blotting of Psd-mNeonGreen-HA (53.5 kDa). MptC, PM-CW marker. **C.** SIM images of log phase cells. Experiments are repeated twice and representative results are shown.

**Table S1.**
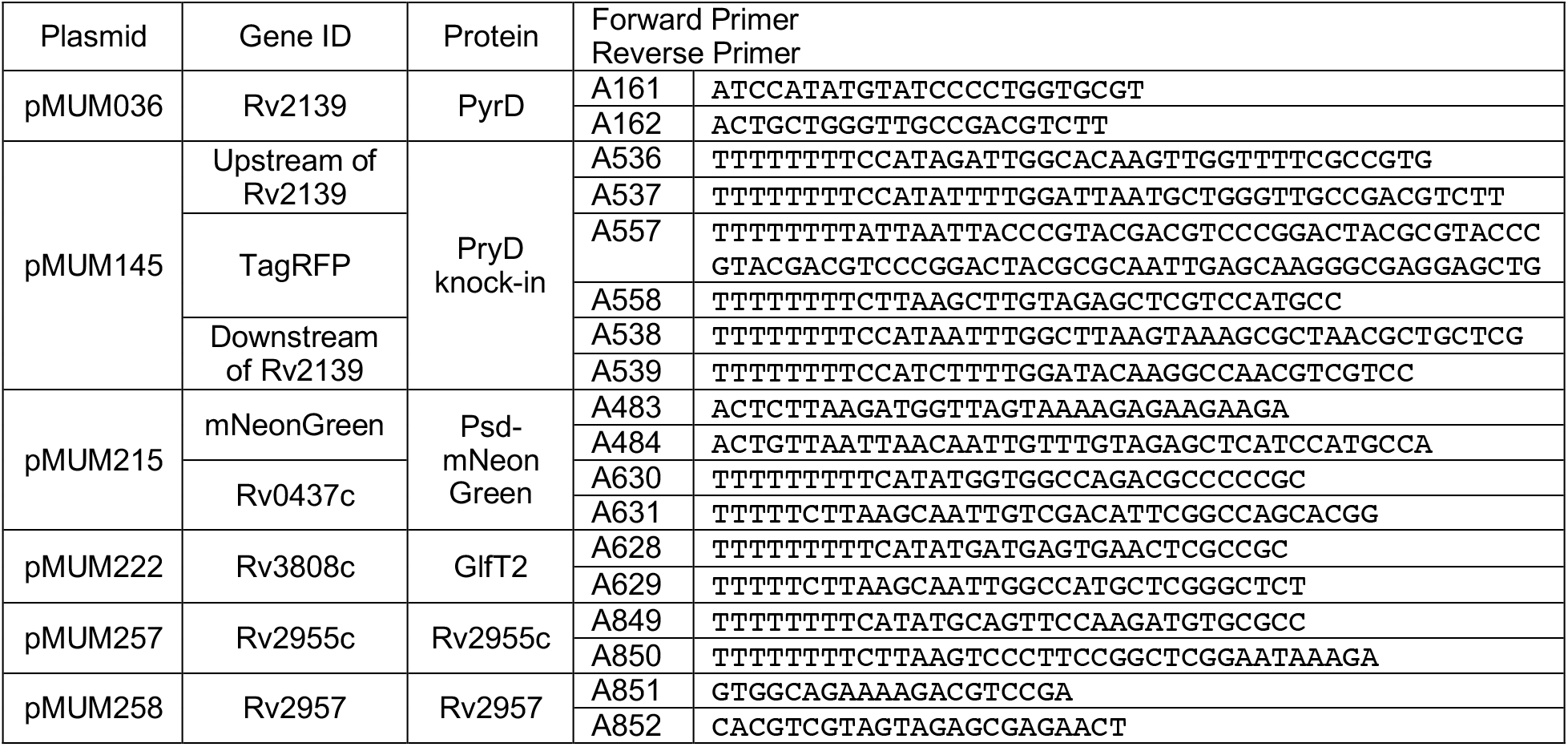
Primers to create plasmids used in this study.

## REFERENCES

1. Kieser KJ, Rubin EJ. 2014. How sisters grow apart: mycobacterial growth and division. Nat Rev Microbiol 12:550–562.

2. Dulberger CL, Rubin EJ, Boutte CC. 2020. The mycobacterial cell envelope - a moving target. Nat Rev Microbiol 18:47–59.

3. Aldridge BB, Fernandez-Suarez M, Heller D, Ambravaneswaran V, Irimia D, Toner M, Fortune SM. 2012. Asymmetry and aging of mycobacterial cells lead to variable growth and antibiotic susceptibility. Science 335:100–104.

4. Rego EH, Audette RE, Rubin EJ. 2017. Deletion of a mycobacterial divisome factor collapses single-cell phenotypic heterogeneity. Nature 546:153–157.

5. García-Heredia A, Pohane AA, Melzer ES, Carr CR, Fiolek TJ, Rundell SR, Chuin Lim H, Wagner JC, Morita YS, Swarts BM, Siegrist MS. 2018. Peptidoglycan precursor synthesis along the sidewall of pole-growing mycobacteria. eLife 7:e37243.

6. Minnikin DE. 1982. Lipids: complex lipids, their chemistry biosynthesis and roles, p. 95–184. In Ratledge, C, Stanford, J (eds.), The Biology of the Mycobacteria: Physiology, Identification and Classification. Academic Press, London.

7. Rahlwes KC, Sparks IL, Morita YS. 2019. Cell walls and membranes of Actinobacteria. Subcell Biochem 92:417–469.

8. Jankute M, Cox JAG, Harrison J, Besra GS. 2015. Assembly of the mycobacterial cell wall. Annu Rev Microbiol 69:405–423.

9. Jackson M. 2014. The mycobacterial cell envelope-lipids. Cold Spring Harb Perspect Med 4:a021105–a021105.

10. Puffal J, García-Heredia A, Rahlwes KC, Siegrist MS, Morita YS. 2018. Spatial control of cell envelope biosynthesis in mycobacteria. Pathog Dis 76:fty027.

11. Meniche X, Otten R, Siegrist MS, Baer CE, Murphy KC, Bertozzi CR, Sassetti CM. 2014. Subpolar addition of new cell wall is directed by DivIVA in mycobacteria. Proc Natl Acad Sci U S A 111:E3243–51.

12. Melzer ES, Sein CE, Chambers JJ, Siegrist MS. 2018. DivIVA concentrates mycobacterial cell envelope assembly for initiation and stabilization of polar growth. Cytoskeleton 75:498–507.

13. Baranowski C, Welsh MA, Sham L-T, Eskandarian HA, Lim HC, Kieser KJ, Wagner JC, McKinney JD, Fantner GE, Ioerger TR, Walker S, Bernhardt TG, Rubin EJ, Rego EH. 2018. Maturing *Mycobacterium smegmatis* peptidoglycan requires non-canonical crosslinks to maintain shape. eLife 7:e37516.

14. Christensen H, Garton NJ, Horobin RW, Minnikin DE, Barer MR. 1999. Lipid domains of mycobacteria studied with fluorescent molecular probes. Mol Microbiol 31:1561–1572.

15. Maloney E, Madiraju SC, Rajagopalan M, Madiraju M. 2011. Localization of acidic phospholipid cardiolipin and DnaA in mycobacteria. Tuberc Edinb Scotl 91 Suppl 1:S150–155.

16. Hayashi JM, Luo C-Y, Mayfield JA, Hsu T, Fukuda T, Walfield AL, Giffen SR, Leszyk JD, Baer CE, Bennion OT, Madduri A, Shaffer SA, Aldridge BB, Sassetti CM, Sandler SJ, Kinoshita T, Moody DB, Morita YS. 2016. Spatially distinct and metabolically active membrane domain in mycobacteria. Proc Natl Acad Sci U S A 113:5400–5405.

17. Morita YS, Velasquez R, Taig E, Waller RF, Patterson JH, Tull D, Williams SJ, Billman-Jacobe H, McConville MJ. 2005. Compartmentalization of lipid biosynthesis in mycobacteria. J Biol Chem 280:21645–21652.

18. Hayashi JM, Richardson K, Melzer ES, Sandler SJ, Aldridge BB, Siegrist MS, Morita YS. 2018. Stress-induced reorganization of the mycobacterial membrane domain. mBio 9:e01823–17.

19. Puffal J, Mayfield JA, Moody DB, Morita YS. 2018. Demethylmenaquinone methyl transferase is a membrane domain-associated protein essential for menaquinone homeostasis in *Mycobacterium smegmatis*. Front Microbiol 9:3145.

20. Plocinski P, Arora N, Sarva K, Blaszczyk E, Qin H, Das N, Plocinska R, Ziolkiewicz M, Dziadek J, Kiran M, Gorla P, Cross TA, Madiraju M, Rajagopalan M. 2012. *Mycobacterium tuberculosis* CwsA interacts with CrgA and Wag31, and the CrgA-CwsA complex is involved in peptidoglycan synthesis and cell shape determination. J Bacteriol 194:6398– 6409.

21. Gola S, Munder T, Casonato S, Manganelli R, Vicente M. 2015. The essential role of SepF in mycobacterial division. Mol Microbiol 97:560–576.

22. Chauhan A, Madiraju MVVS, Fol M, Lofton H, Maloney E, Reynolds R, Rajagopalan M. 2006. *Mycobacterium tuberculosis* cells growing in macrophages are filamentous and deficient in FtsZ rings. J Bacteriol 188:1856–1865.

23. Plocinska R, Purushotham G, Sarva K, Vadrevu IS, Pandeeti EVP, Arora N, Plocinski P, Madiraju MV, Rajagopalan M. 2012. Septal localization of the *Mycobacterium tuberculosis* MtrB sensor kinase promotes MtrA regulon expression. J Biol Chem 287:23887–23899.

24. Bachhawat N, Mande SC. 1999. Identification of the INO1 gene of *Mycobacterium tuberculosis* H37Rv reveals a novel class of inositol-1-phosphate synthase enzyme. J Mol Biol 291:531–536.

25. Wolucka BA, de Hoffmann E. 1998. Isolation and characterization of the major form of polyprenyl-phospho-mannose from *Mycobacterium smegmatis*. Glycobiology 8:955–962.

26. Takayama K, Schnoes HK, Semmler EJ. 1973. Characterization of the alkali-stable mannophospholipids of *Mycobacterium smegmatis*. Biochim Biophys Acta 316:212–221.

27. Crick DC, Schulbach MC, Zink EE, Macchia M, Barontini S, Besra GS, Brennan PJ. 2000. Polyprenyl phosphate biosynthesis in *Mycobacterium tuberculosis* and *Mycobacterium smegmatis*. J Bacteriol 182:5771–5778.

28. Takayama K, Goldman DS. 1970. Enzymatic synthesis of mannosyl-1-phosphoryl- decaprenol by a cell-free system of *Mycobacterium tuberculosis*. J Biol Chem 245:6251– 6257.

29. Schulbach MC, Brennan PJ, Crick DC. 2000. Identification of a short (C_15_) chain Z- isoprenyl diphosphate synthase and a homologous long (C_50_) chain isoprenyl diphosphate synthase in *Mycobacterium tuberculosis*. J Biol Chem 275:22876–22881.

30. Björnberg O, Rowland P, Larsen S, Jensen KF. 1997. Active site of dihydroorotate dehydrogenase A from *Lactococcus lactis* investigated by chemical modification and mutagenesis. Biochemistry 36:16197–16205.

31. Munier-Lehmann H, Vidalain P-O, Tangy F, Janin YL. 2013. On dihydroorotate dehydrogenases and their inhibitors and uses. J Med Chem 56:3148–3167.

32. García-Heredia A, Kado T, Sein CE, Puffal J, Osman SH, Judd J, Gray TA, Morita YS, Siegrist MS. 2021. Membrane-partitioned cell wall synthesis in mycobacteria. eLife 10:e60263.

33. Onwueme KC, Vos CJ, Zurita J, Soll CE, Quadri LEN. 2005. Identification of phthiodiolone ketoreductase, an enzyme required for production of mycobacterial diacyl phthiocerol virulence factors. J Bacteriol 187:4760–4766.

34. Siméone R, Constant P, Malaga W, Guilhot C, Daffé M, Chalut C. 2007. Molecular dissection of the biosynthetic relationship between phthiocerol and phthiodiolone dimycocerosates and their critical role in the virulence and permeability of *Mycobacterium tuberculosis*. FEBS J 274:1957–1969.

35. Purwantini E, Daniels L, Mukhopadhyay B. 2016. F_420_H_2_ is required for phthiocerol dimycocerosate synthesis in mycobacteria. J Bacteriol 198:2020–2028.

36. Ng V, Zanazzi G, Timpl R, Talts JF, Salzer JL, Brennan PJ, Rambukkana A. 2000. Role of the cell wall phenolic glycolipid-1 in the peripheral nerve predilection of *Mycobacterium leprae*. Cell 103:511–524.

37. Tsenova L, Ellison E, Harbacheuski R, Moreira AL, Kurepina N, Reed MB, Mathema B, Barry CE, Kaplan G. 2005. Virulence of selected *Mycobacterium tuberculosis* clinical isolates in the rabbit model of meningitis is dependent on phenolic glycolipid produced by the bacilli. J Infect Dis 192:98–106.

38. Sinsimer D, Huet G, Manca C, Tsenova L, Koo M-S, Kurepina N, Kana B, Mathema B, Marras SAE, Kreiswirth BN, Guilhot C, Kaplan G. 2008. The phenolic glycolipid of *Mycobacterium tuberculosis* differentially modulates the early host cytokine response but does not in itself confer hypervirulence. Infect Immun 76:3027–3036.

39. Reed MB, Domenech P, Manca C, Su H, Barczak AK, Kreiswirth BN, Kaplan G, Barry CE. 2004. A glycolipid of hypervirulent tuberculosis strains that inhibits the innate immune response. Nature 431:84–87.

40. Camacho LR, Ensergueix D, Perez E, Gicquel B, Guilhot C. 1999. Identification of a virulence gene cluster of *Mycobacterium tuberculosis* by signature-tagged transposon mutagenesis. Mol Microbiol 34:257–267.

41. Baulard AR, Gurcha SS, Engohang-Ndong J, Gouffi K, Locht C, Besra GS. 2003. *In vivo* interaction between the polyprenol phosphate mannose synthase Ppm1 and the integral membrane protein Ppm2 from *Mycobacterium smegmatis* revealed by a bacterial two-hybrid system. J Biol Chem 278:2242–2248.

42. Rokicki CAZ, Brenner JR, Dills AH, Judd JJ, Kester JC, Puffal J, Sparks IL, Prithviraj M, Anderson BR, Wade JT, Gray TA, Derbyshire KM, Fortune SM, Morita YS. 2021. Fluorescence imaging-based discovery of membrane domain-associated proteins in *Mycobacterium smegmatis*. J Bacteriol 203:e0041921.

43. Malhotra S, Vedithi SC, Blundell TL. 2017. Decoding the similarities and differences among mycobacterial species. PLoS Negl Trop Dis 11:e0005883.

44. Namouchi A, Cimino M, Favre-Rochex S, Charles P, Gicquel B. 2017. Phenotypic and genomic comparison of *Mycobacterium aurum* and surrogate model species to *Mycobacterium tuberculosis*: implications for drug discovery. BMC Genomics 18:530.

45. Onwueme KC, Vos CJ, Zurita J, Soll CE, Quadri LEN. 2005. Identification of phthiodiolone ketoreductase, an enzyme required for production of mycobacterial diacyl phthiocerol virulence factors. J Bacteriol 187:4760–4766.

46. Pérez E, Constant P, Laval F, Lemassu A, Lanéelle M-A, Daffé M, Guilhot C. 2004. Molecular dissection of the role of two methyltransferases in the biosynthesis of phenolglycolipids and phthiocerol dimycoserosate in the *Mycobacterium tuberculosis* complex. J Biol Chem 279:42584–42592.

47. Fukuda T, Matsumura T, Ato M, Hamasaki M, Nishiuchi Y, Murakami Y, Maeda Y, Yoshimori T, Matsumoto S, Kobayashi K, Kinoshita T, Morita YS. 2013. Critical roles for lipomannan and lipoarabinomannan in cell wall integrity of mycobacteria and pathogenesis of tuberculosis. mBio 4:e00472–12.

48. Paintdakhi A, Parry B, Campos M, Irnov I, Elf J, Surovtsev I, Jacobs-Wagner C. 2016. Oufti: an integrated software package for high-accuracy, high-throughput quantitative microscopy analysis. Mol Microbiol 99:767–777.

